# An anomalous 3’-terminal phosphorothioated mismatch bypass activity and its application as a binary molecular switch

**DOI:** 10.1101/2024.07.27.605420

**Authors:** Shrawan Kumar, Hira Singh Gariya, Chandrika Sharma, Saba Parveen, Vandana K Nair, Mrittika Sengupta, Souradyuti Ghosh

## Abstract

Phosphorothioated (PST) oligonucleotides are increasingly being used in RNA silencing, antisense, and biosensing applications. However, the possibilities and consequences of their desultory interactions with other possible nucleic acids and DNA polymerases inside the cell remain inadequately characterized. In this study, we report the discovery of an unusual terminal mismatch bypass activity involving 3′-PST containing DNA primers and certain strand displacement DNA polymerases. Using rolling circle DNA amplification, we have identified that strand displacement DNA polymerases such as phi29 and BST large fragment (LF) can bypass 3′-terminal PST mismatches upto 1 – 20 nt length. Next, we explore the length and sequence dependence of this unusual attribute, incubation in near-ambient and 60 – 65°C temperatures, and measures to blockade or modulate this mismatch bypass activity to create a binary fully nucleic acid-based and non-photocontrolled molecular switch (the first of its kind). After proposing possible underlying mechanisms for this activity, we discuss its potential consequences and applications.

## Introduction

DNA polymerases utilize their exonuclease activity, duplex structural distortion, and several auxiliary protein complexes to discriminate between correct vs incorrect base pairing during dNTP incorporation^1,2^. As even a single mismatch could lead to biologically detrimental consequences, polymerases are equipped with a proofreading mechanism to recognize the mismatch presence via the thermodynamic stability of the primer-template complex. The unfavorable alignment of the mismatched bases would affect the optimum distance for catalytic nucleotide incorporation efficiency, lowering and ultimately stalling the rate of nucleic acid polymerization^3,4^. Exonuclease activity in the polymerases further bolsters the mismatch recognition process.

Chemical modifications such as PST linkages in oligonucleotides would confer stability against various nucleases and polymerase exonuclease properties and, therefore, have seen extensive utilization in antisense and siRNA technologies^5–7^. Additionally, the extent of phosphorothioate content may modulate the cellular uptake and extrahepatic distribution, therefore regulating the therapeutic efficacy^8–10^. However, it also alters the intracellular interaction of the oligonucleotide in terms of protein binding, H-bonding, as well as delivery pathways^11,12^. As a result, it may affect how polymerases process such linkages. Interestingly, despite the frequent application of PST linkage-containing oligonucleotides in nucleic acid therapeutics, their processing or off-target effects by various DNA polymerases inside or outside the target cell types have not been closely studied. Such PST oligonucleotides may partially bind genomic DNA, circular mitochondrial DNA^13^, or extrachromosomal circular DNA^14^, the last being a recently identified hallmark of several cancers. In such cases, the partially annealed region with chemically modified nuclease-resistant non-complementary 3′-region may appear as a primer with an extended terminal mismatch for a DNA or RNA polymerase.

In prior studies, the efficiency of single base mismatch extensions by prokaryotic proofreading-lacking strand displacement polymerases (BST LF, klenow 3′→5′ exo-(klenow exo-), and vent exo(-)) were higher for 3′-single PST primer than unmodified phosphodiester primer^15^. In comparison, aside from the inability of phi29 (another strand displacement polymerase with exonuclease ability towards phosphodiester 3′-mismatches and 3′-overhangs^16,17^) to digest PST oligonucleotides, their interaction remains uninvestigated. In eukaryotes, DNA polymerase δ is the lagging strand displacement polymerase and a critical partner in strand displacement activity during homologous recognition. It could remove or bypass phosphodiester-containing 3′-terminal single mismatches in the presence or absence of its exonuclease activity, respectively^18^. However, it remains unexplored whether it could bypass 3′-single nucleotide or longer 3′-terminal mismatches having PST linkages. Similarly, the phosphorothioated mismatch processing ability of the strand displacing mitochondrial DNA polymerase (DNA pol γ_19,20_) is unexplored. Due to the increasing utilization of PST oligonucleotides in therapeutics and diagnostics, probing their interaction with mismatch-bypassing polymerases and endogenous nucleic acids would be of critical interest.

In this study, we report our preliminary findings of a unique, sequence-independent, and unusual 3′-terminal mismatch bypass ability of strand displacement polymerases for 3′-triple PST linkage present in DNA oligonucleotides. Due to the ample in vivo presence of various endogenous circular DNA substrates and strand displacement polymerases, we utilized rolling circle amplification as an equivalent setup (i.e., using a circular DNA (cDNA) template, a PST primer, and a strand displacement polymerase) to probe this observation. We have investigated combinations of cDNA with PST primers with mismatches starting from 1 to 11, 16, or 20 nt, and three types of DNA polymerases. Both identical (e.g., A-A, G-G) and non-identical mismatches (e.g., A-G and T-C) were studied. Finally, we also demonstrated a method for modulating the terminal mismatch bypass activity and, in principle, its potential utilization in biocomputing and biosensing research.

## Materials and Methods

The oligonucleotides used in this study (listed in Table S1) were purchased from Eurofins Genomics or Sigma Aldrich after the supplier provided high purity salt-free (HPSF, for Eurofin) or desalting (for Sigma) purification method and used without further purification. T4 polynucleotide kinase (PNK), T4 DNA ligase, phi29 DNA polymerase, klenow exo-, BST large fragment (LF) polymerase, dNTP, BSA, MgSO_4_, isothermal amplification buffer, and phi29 buffer were purchased from New England Biolabs, USA. Tris buffers, NaCl, spermidine, and dithioerythritol (DTT) were purchased from Sisco Research Laboratories Pvt. Ltd. (SRL) -India. SYBR™ Green I Nucleic Acid Gel Stain 10,000X was purchased from Invitrogen. All solutions were prepared using ultrapure water obtained from a Millipore Milli-Q Type II water purification system.

### 5′-Phosphorylation

Stock concentrations (100 μM) of linear ssDNA oligonucleotides that are precursor to cDNA were diluted with DEPC treated water to final concentration 16 μM. The solutions were then snap-cooled by 5 minutes heating at 95°C followed by 5 minutes of incubation on ice to linearize the DNA. The 5′-phosphorylation was carried on the oligonucleotides (final concentration 8 μM) in presence of T4 PNK (0.375 Units L^-1^), ATP (final 1 mM), NEB ligase buffer (final 50 mM Tris-HCl pH 7.5, 10 mM MgCl_2_, 1 mM ATP, 10 mM DTT), DTT (final 5 mM), and spermidine (final 1.7 mM), where all reagents were added with to the tube on ice, followed by incubation at 37°C for 3 h. For self-annealing CD3, the solution was then subjected to annealing (95°C to 4°C, with simultaneous inactivation of T4 PNK). For splint-padlock ligation (CD1.Xm and CD2.Xm series), the phosphorylation mixture containing the padlock oligonucleotide was first heat inactivated (85°C 20 min). Next, splint (final concentration 7.4 μM) was mixed with a 5′-phosphorylated padlock precursor (prepared as described above, final concentration 6.1 μM), and the solution was subjected to aforementioned annealing step.

### Ligation

Circularization of the annealed 5’-phosphorylated DNA (final concentration 6.1 μM for precursors to CD1.Xm and CD2.Xm series and 3.2 μM for CD3) was carried out in presence of T4 DNA ligase (8 Cohesive End Units μL^-1^), ATP (final 1 mM), and T4 DNA ligase buffer (final 50 mM Tris-HCl pH 7.5, 10 mM MgCl_2_, 1 mM ATP, 10 mM DTT), where the requisite reagents were added with the tube on ice. The ligation was carried out by incubating the mixture at 16°C for 16 hours and then enzyme inactivation at 75°C for 20 min. The ligation was confirmed by running the reaction in 13% denaturing PAGE. Oligonucleotides of known concentrations were run side by side in the gel to assess the concentrations of the ligated oligonucleotides.

### Exonucleases treatment

Following ligation, the reaction mixture was subjected to snap-cooling. Next, exonuclease I (final 2.4 units μL^-1^) and III (final 12 units µL^-1^) were added in exonuclease III buffer (final 10 mM Bis-Tris-Propane-HCl pH 7, 10 mM MgCl_2_, 1 mM DTT) while the tube was kept in ice. The reaction mixture was then incubated at 37°C for 4 h followed by enzyme inactivation at 85°C for 20 minutes and stored in -20 °C for future use. The reaction was carried out in 25 μL reaction volume with final oligonucleotide concentrations of 1.5 μM for both self-annealed and splint-padlock ligation substrates.

### Real time RCA

The above circular DNA was annealed with respective primers (and blockers, when applicable) at 30°C for 30 min in 50 mM Tris-HCl (pH 8) and 50 mM NaCl. The real-time RCA was performed in 15 μL volume containing circular DNA (final 0.02 μM), primer (final 0.02 μM), blocker (when applicable, final 0.02 μM), dNTPs (final 1.0 mM), BSA (final 0.2 μg uL^-1^), phi29 buffer (final 50 mM Tris-HCl pH 7.5, 10 mM MgCl_2_, 10 mM (NH_4_)_2_SO_4_, 4 mM DTT), 0.4X SYBR Green I (final concentration), and phi29 DNA polymerase (final 0.2 units μL^-1^) or klenow exo-enzyme (final 1 unit μL^-1^). For klenow exo-enzyme, 1X NEB 2 buffer (final 50 mM NaCl, 10 mM Tris-HCl, 10 mM MgCl_2_, 1 mM DTT, pH 7.9 @ 25°C) was used. Then RCA reaction was performed at 30°C (or 37°C for klenow exo(-)) for 2 hours and fluorescence signals were recorded at intervals of 2 min. For BST LF polymerase assisted amplification, a typical 15 μL reaction involved BST LF polymerase (final 1.6 units μL^−1^), isothermal amplification buffer (final 20 mM Tris-HCl, 10 mM (NH_4_)_2_SO_4_, 50 mM KCl, 2 mM MgSO_4_, 0.1% Tween® 20, pH 8.8@25°C), MgSO_4_ (final 4 mM), dNTP (final 1.4 mM), SYBR Green I (final 0.6X). The reaction involved incubation at 55°C and fluorescence signals were recorded at intervals of 2 min.

### Baseline correction methods and end-RFU measurement

In this study, any software-imposed baseline corrections and subtractions were removed from the amplification profiles and absolute amplification profiles were generated. Next, a manual subtraction of the 1^st^ cycle RFU magnitude of “cDNA alone” control (present in all experiments) in each assay from the rest of the amplification profiles was performed to bring uniformity. The end-RFUs were acquired from this manual-baseline-subtracted amplification profiles.

## Results and Discussion

### Design of oligonucleotides, experimental settings, and data analysis

The DNA oligonucleotides used in this study were named by the following convention and listed in Table S1. We utilized a common triple 3′-PST primer and a non-PST primer for all studies, with identical sequence and length (41 nt). At the same time, the mismatch sequence region and length of the same were modified in the cDNA itself. The cDNA series having identical (i.e., A-A, G-G, C-C, and T-T) or non-identical (i.e., A-G, T-C) series of mismatches was denoted as CD1.Xm and CD2.Xm, respectively. Here, X represents the length of the mismatch region, while m represents the mismatch. For instance, CD2.5m would suggest a cDNA with 5 terminal non-identical mismatches. Additionally, a cDNA CD_t-match, providing a 6 nt internal mismatch and a 5 nt terminal match, was also incorporated as a control to the study.

### Measurement of amplification

The oligonucleotide primers (primer_PST or primer_nonPST) and cDNA containing the mismatch were subjected to phi29-mediated real-time RCA for end-fluorescence measurement. The terminal mismatch bypass activity has been presented by normalized end-RFU for X nt mismatches (X = 0 – 11, 16, 20 nt). It represented the ratio of end-fluorescence between an X nt long mismatch and that without any mismatch (i.e., from CD1.0m). For both the primer_PST and primer_nonPST, ideally, the 0 nt mismatch should produce fluorescence and thus have a normalized end-RFU of 1. For primer_nonPST forming any non-zero nt length mismatches with the cDNA, including that with a terminal match, various degrees of non-zero (and presumably close to 1) end-RFU would be anticipated. This would stem from the 3′-exonuclease activity of phi29 DNA polymerase digesting any phosphodiester backbone mismatches^17^. On the other hand, if terminal bypass of a 3′-mismatch for phosphorothioate primer is not occurring, the normalized end-RFU of all mismatches except 0 nt and terminal mismatch should be zero.

For the 1 – 11 nt non-PST mismatches and the terminal match, the normalized end-RFU was expectedly detected at various non-zero magnitudes (Figure 1A – C and Figure S1). This observation corroborated earlier studies and was due to the 3′-exonuclease activity of the phi29 polymerase^17,21^. Interestingly, when 1 – 11 nt identical mismatches were present against the primer_PST, it unexpectedly led to “unusual” non-zero normalized end-RFU (Figure 1C and Figure S2). We initially speculated that this could be due to an HRCA-like mechanism where small amounts of amplicons could be non-specifically generated by phi29-mediated non-specific RCA (Scheme S1). These amplicons would have complementarity to the cDNA and hence act as an RCA primer. To validate or vindicate this speculation, we designed a series of cDNAs having 1 – 11 nt non-identical mismatches (i.e., A-G, T-C) against the primer_PST (CD2.Xm series, Table S1). Similar to identical mismatch containing studies (above), both PST and non-PST primers were investigated. Once again, all of the 1 – 11 nt mismatches against primer_PST led to unexpected non-zero normalized end-RFU while primer_nonPST produced anticipated non-zero fluorescence to various degrees (Figure 1D, Figure S3 and S4). A minor enhancement of amplification at 1, 6, or 7 nt, and then again towards 11 nt PST mismatch was noticeable at both sets of experiments (identical as well as non-identical mismatch). The similar magnitudes of normalized end-RFU for the non-identical mismatch containing primer_PST indicated the minimal contribution of the aforementioned HRCA-like mechanism. Terminal bypass of non-identical PST mismatches was found for 16 and 20 nt long mismatches (Figure S5). The current study did not investigate whether this bypass pathway existed for mismatches longer than 20 nt.

**Figure 1.**
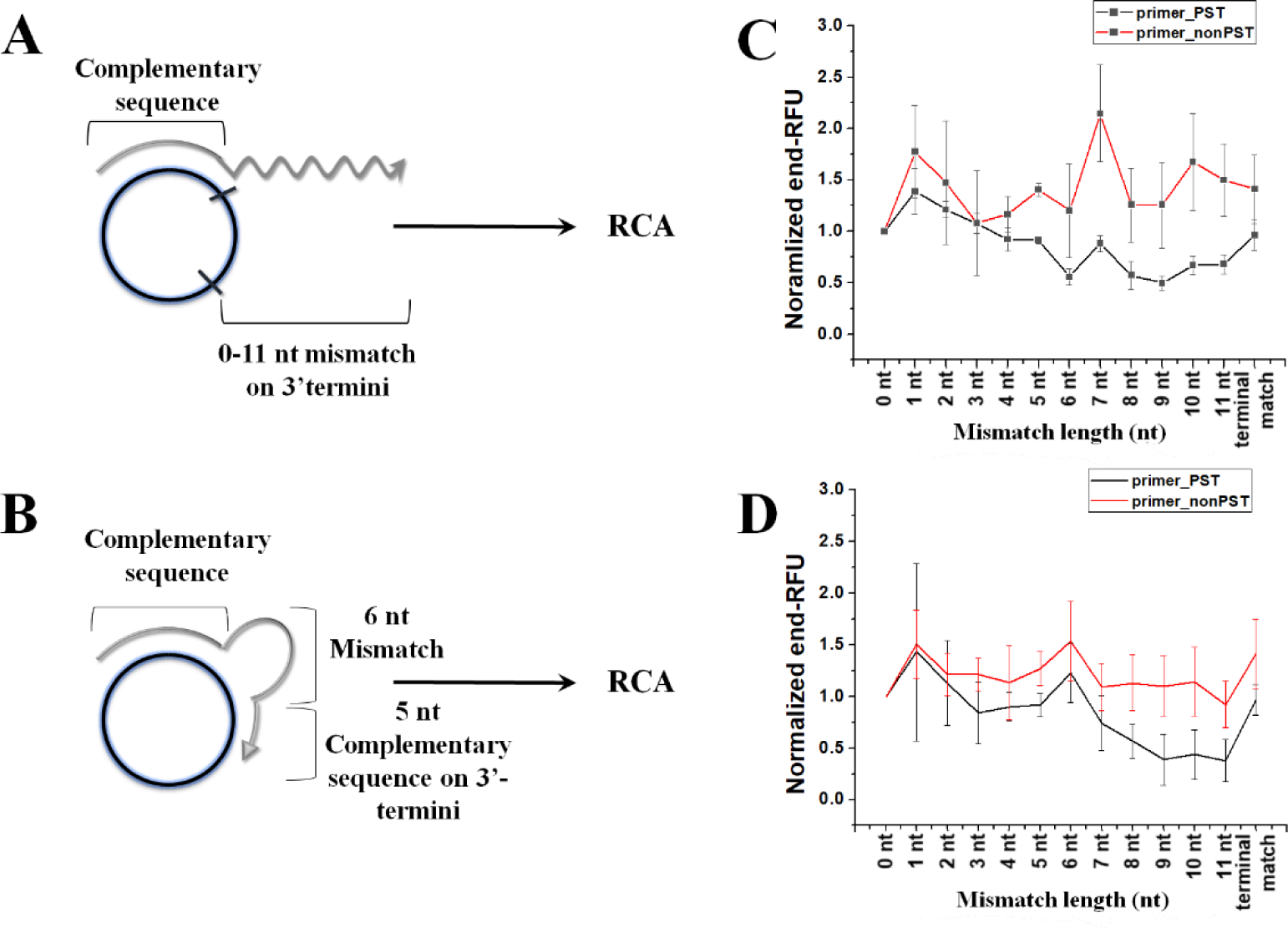
Quantification of 3′-terminal mismatch bypass for a cDNA template and phosphorothioated primer. A and B, Schematics of binding between cDNA template and primers (primer_PST and primer_nonPST) having 0 – 11 nt mismatches or terminal matches, respectively. C and D, Normalized end-RFU for identical or non-identical 0 – 11 nt mismatches, respectively.

### Terminal mismatch bypass using other phosphorothioated sequences and polymerases

Next, we sought to probe whether the observed terminal mismatch bypass could be a primer-specific observation, as all the above experiments contained only a single PST primer with various cDNAs. We also wanted to investigate if any strand displacement polymerase other than phi29 may demonstrate the aforementioned terminal mismatch behavior. To study the former, we utilized another set of phosphorothioated primers PST3.0m (having 0 nt mismatch), PST3.1m (1 nt mismatch), PST3.2m (2 nt mismatch) against cDNA CD3 (sequences in Table S1). The corresponding RCA experiment used phi29 and klenow exo-DNA polymerases (Figure S6D and E, respectively). RCA with 0 – 2 nt mismatches using phi29 again demonstrated amplification. On the other hand, RCA involving klenow exo-(which cannot distinguish between PST and non-PST primer) recorded similar amplification signals for 1 nt mismatch but little or none for the 2 nt. The latter observation was consistent with earlier single terminal mismatch bypass activity by klenow exo-for a short unmodified primer^22^.

BST LF polymerase is a thermophilic DNA polymerase lacking exonuclease activity and is extensively used in loop-mediated isothermal amplification. Without an exonuclease proofreading activity, a 3′-phosphorothioate or 3′-phosphodiester-based primer would appear identical to it except for any differential biophysical interaction between the primers and the polymerase active site. Treatment of cDNAs having non-identical mismatches (CD2.Xm series) with primer_PST and primer_nonPST in the presence of BST LF polymerase was performed to evaluate the possibility of terminal mismatch bypass (Figure S7 and S8). For both the primer_PST and primer_nonPST, terminal mismatch bypass activity was demonstrated until 5 nt long mismatch. Interestingly, higher amplification efficiency was present for mismatches than without any mismatch in both cases. Surprisingly, both did not show any amplification for the 4 nt long mismatches. Despite a yet unknown mechanism, these findings demonstrated the terminal mismatch bypass activity to be sequence-independent and occurring for other strand displacement polymerases to a varying degree of mismatch length.

### Modulating terminal-mismatch bypassing amplification using a blocker

Next, we explored whether the PST-based terminal mismatch could be somehow modulated. We hypothesized that the single-stranded 3′-terminal PST region was somehow responsible for either directly or indirectly (for instance, via its amplification-generated complement) annealing to the cDNA (see below for possible mechanisms). Therefore, a sequence binding the single-stranded mismatched region might, in principle, “block” this terminal mismatch bypass activity. To explore further, we designed two such “blocker” sequences (blocker_16 and blocker_20, T_m_ 44.5°C and 51.4°C, respectively, calculated for 0.25 μM oligonucleotide at 50 mM Na^+^) for 16 and 20 nt non-identical mismatches. These oligonucleotides would be utilized to completely stop or at least suppress the amplification between the respective cDNA-primer pairs (i.e., cDNAs CD2.16m or CD2.20m with their PST or non-PST primers). Indeed, amplification of the cDNA-mismatched primer pairs in the presence of both 16 and 20 nt blockers demonstrated that it could be attenuated (Figure 2B and C, Figure S9C – F). For amplification concerning cDNA (both CD2.16m and CD2.20m) and primer_nonPST, the end-RFUs were independent of the presence or absence of the respective blocker sequences. The latter originated from the 3′-exonuclease activity of phi29 polymerase digesting both the primer and the blocker, thus creating a suitable primer for RCA^17^. Interesringly, the blocker could not be removed despite its phosphodiester backbone and subsequence susceptibility to phi29 3′-exonuclease property. The observation that 20 nt blocker is relatively more effective in suppressing could be owing to its greater thermodynamic stable binding with the primer_PST. So, the presence or absence of the blocker here acted as an input to generate binary “Off” or “On’’ output switch, similar to a single “bit” in computing (Figure 2D). In that respect, the blocker’s role is analogous to that of a resistor in a current-transmitting electrical circuit. To the best of our knowledge, this would be the first example of a fully nucleic acid-based and non-photocontrolled molecular switch that could be utilized to “turn on” or “turn off” polymerase-assisted amplification.

**Figure 2.**
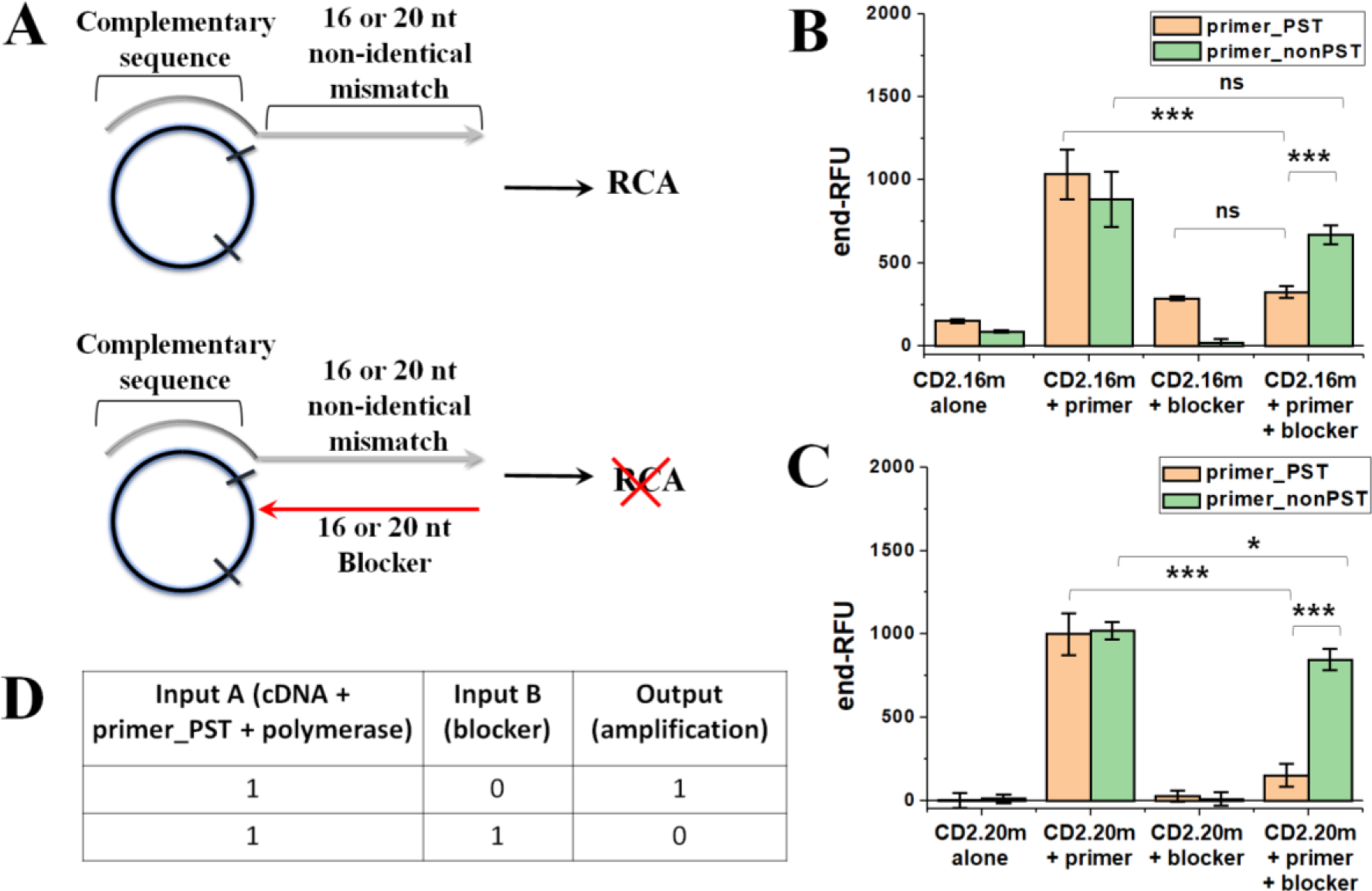
Effect of “blocker” sequence on terminal mismatch bypass amplification. A, Schematics of “turning on/off” terminal mismatch bypass amplification using a 16 or 20 nt “blocker” sequence. B and C, Regulation of terminal mismatch bypass amplification for 16 and 20 nt mismatch-blocker combinations, respectively. D, Truth table for binary logic gate. Phi29 DNA polymerase was used in this experiment. *P ≤ 0.05, **P ≤ 0.01, ***P ≤ 0.001.

### Possible mechanisms for terminal mismatch bypass and future applications

The above experiments, including ones involving the blocker sequences, conclusively demonstrated that the unusual terminal mismatch bypass activity stemmed from the phosphorothioated mismatch region. To rationalize this novel but unusual terminal mismatch bypass activity, we first hypothesized that it could originate from an intramolecular “phosphorothioated-terminal hairpin formation” followed by “self-priming extension”. This mechanism could be further justified from studies by Prof Ellington and co-workers demonstrating such amplification at both 60-65°Cand near ambient temperature (40°C)^23,24^. However, we argue the possible non-applicability of this mechanism here due to the following reasons. NUPACK simulations did not show a terminal hairpin formation (a necessity for self-priming amplification) in the PST primer (i.e., primer_PST) used in this study, even at high salt concentrations (Figure S10)^25^. Secondly, a terminal hairpin formation would not be possible for shorter mismatches (1 – 5 nt) when the primer_PST is annealed to the cDNA. However, amplifications were recorded for 1 – 5 nt mismatches for both the phi29 and BST LF polymerases. Additionally, any amplicon generated from self-priming would preferably bind to itself rather than the cDNA due to entropic reasons. Such a mechanism thus may not provide high magnitudes of end-RFU, as seen here. Finally, self-priming amplification would have occurred for 1 – 20 nt mismatches for both phi29 and BST LF DNA polymerases. However, contrary to this anticipation, no such amplifications were recorded beyond 5 nt mismatch for either primer_PST or primer_nonPST in case of the BST LF DNA polymerase. The last observation probably indicated that the underlying mechanism could involve a structure that may undergo a temperature-dependent destabilization at higher temperature, therefore rationalizing lack of amplifications for longer mismatches.

Instead, we propose a proximity-dependent transient intermolecular annealing to elucidate the above observation (Scheme 1A). This relies on lipophilic nature of phosphorothioate linkages and its base stacking as well as hydrophobic interactions with aromatic amino acids such as tyrosine, several of which are present at the phi29 active site^26,27^. In this mechanism, instead of forming a self-priming hairpin loop, the mismatch region non-sequence-specifically and temporarily binds to cDNA simply due to proximity and relatively higher hydrophobicity of the phosphorothioate linkage. This association leads to the formation of a transient primer-cDNA template-like structure that is recognized by DNA polymerase, which “latches” onto it and causes amplification. This hypothesis would rationalize the similar degree of normalized end-RFU for both the identical and non-identical mismatches (Figure 1C and D). This hypothesis could also be supported by the absence of amplification beyond the 5 nt for BST LF polymerase, caused by higher temperature disrupting the transient intermolecular complexation between primer_PST and cDNAs.

In the case of therapeutic applications of PST oligonucleotides, such unintended amplifications may have severe clinical consequences. For e.g., amplicons generated from terminal mismatch bypass amplifications between PST oligonucleotides and endogenous oncogenic extrachromosomal cDNAs would lead to multiple gene copies present in the cell. Similarly, unwanted amplifications may cause the generation of otherwise highly regulated mobile genetic elements, leading to unpredictable mutations. Therefore, interactions between PST oligonucleotides and various genetic elements beyond their intended target may warrant thorough genomewide probing. Future work from our lab would study the correlation between the degree of PST modification in the oligonucleotide backbone and the terminal mismatch bypass.

**Scheme 1.**
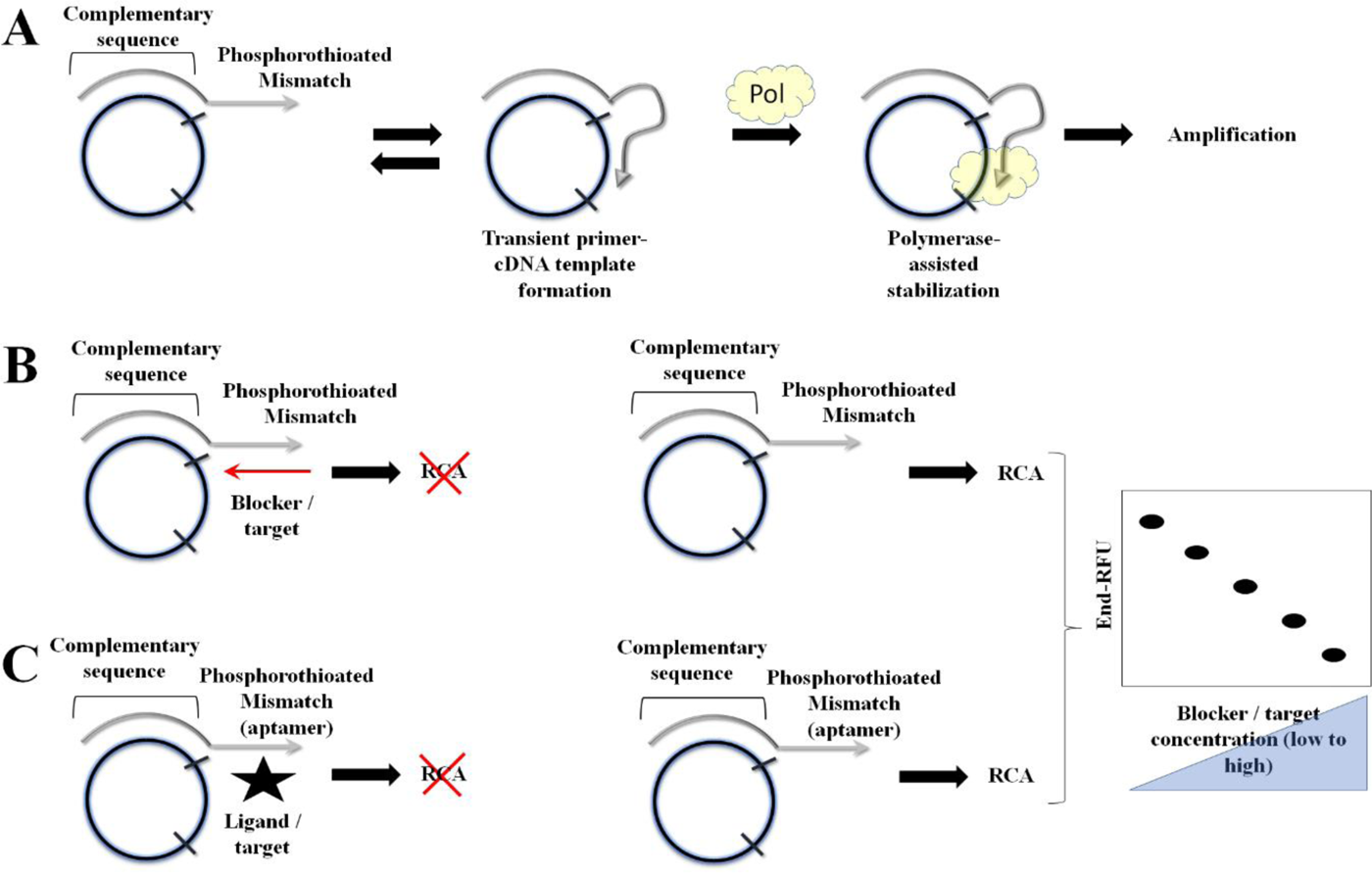
A, Proposed mechanism for terminal mismatch bypass amplification. B and C, Possible utilization of terminal mismatch bypass amplification for nucleic acid and aptameric target sensing, respectively.

Several possible applications of this terminal mismatch bypass activity could be envisioned, especially in biocomputation and biosensing. As explained earlier, it can be used as a binary molecular switch system (Figure 2D). Multi-input molecular switches can be created through the incorporation of multiple fluorescently tagged molecular-beacon-like reporter sequences responsive to the RCA amplicons. Due to its sequence-independent nature, the system can be designed to recognize a nucleic acid sequence as a target (Scheme 1B). We also anticipate that if an aptamer is present within the mismatch region, a similar ligand-recognizing “on/off” switch can be constructed (Scheme 1C). In either case, when the targets (nucleic acid or aptameric ligand) are absent, the molecular switch would “turn on”. The presence of the analyte (a nucleic acid in Scheme 1B and ligand/target in Scheme 1C) in equal concentration as that of mismatch-cDNA assembly would then cause the molecular switch to completely “turn off” (as in Figure 2C). There would presumably be multiple intermediate partial “off” states (having non-zero amplification magnitude) between the two extreme “on/off” scenarios. We anticipate that the fluorescence difference between the “off”, “partial off”, and “on” states could be correlated with analyte concentrations, thus creating an effective functional nucleic acid biosensing platform.

A therapeutic application aimed at controlled silencing or modulating gene expression could be envisioned where a pre-annealed PST primer-circular DNA pair would be transfected to the cell, with the circular DNA carrying gene regulating elements such as a DNAzyme in it. The pair would be stable in biological environment thanks to both circular DNA and the PST primer being nuclease resistant. Once inside nucleus or mitochondrial matrix, both of which have strand displacement polymerases, it will trigger release of multiple copies of DNAzyme, targeting native RNA molecule and modulating gene expression. When the phosphorothioated primer would carry a 3′-mistamch region (as in this case), a subsequent addition of a complementary “blocker” would stop the polymerase activity, and resuming native gene expression level. These experiments will be attempted in future studies from our lab.

## Conclusion

In summary, this study reports the discovery of a novel 3′-terminal mismatch bypass property for PST oligonucleotides by strand displacement polymerases using the RCA platform as a setup. This attribute was sequence-independent, observed in identical as well as non-identical mismatches, and could be seen at least upto 20 nt mismatch length. We also demonstrate that the observed bypass activity could be “turned off” using an additional “resistor”-like blocker sequence, creating a binary switch. We hypothesize that this could originate from transient interaction between PST 3′-termini of the primer, cDNA template, and DNA polymerase. This discovery recommends a closer look at the interaction between PST oligonucleotides and various endogenously present circular nucleic acid structures, more so due to the former’s increasing involvement in molecular therapeutics. This discovery is also anticipated to lead to novel applications in the biosensing and biocomputing domain.

## Author Contributions

SK, HSG, and SG designed the study. SK, HSG, CS, and SP performed the experiments. MS provided inputs regarding DNA repair and molecular biology aspects of strand displacement DNA polymerases. SK, HSG, and SG performed data analyses and wrote the manuscript. All authors edited and approved the final manuscript.

## Notes

The research team has filed an Indian patent application (filing no 202341080341) related to this discovery.

## Acknowledgments

This work was supported by the Early Career Research Grant (No. ECR/20l8/001399) awarded by the Department of Science and Technology Science and Engineering Research Board, India, and Mahindra University internal funding. The authors thank lab managers Mr. Saddam Abdul Aziz Khan and Mr Rajat Tiwari for their assistance in day-to-day lab management. The authors are grateful to Prof. Rajinder S. Chauhan for his constant scholarly support.

## Supporting Information

Oligonucleotide sequences, representative amplification plots, and additional molecular mechanisms.

