## Supplementary Information for "An anomalous 3’-terminal phosphorothioated mismatch bypass activity and its application as a binary molecular switch"

**Table S1.** Sequences and description of oligonucleotides used in this study. Splint binding region: underlined; Primer binding region: italic; Mismatch region: in red font.

| Oligonucleotide Name | Sequence (5′ → 3′) | Description |
|---|---|---|
| **Splint** | GGTCGCCTC CTAGGATC | Common splint for all circular DNA (cDNA) |
| **CD1.0m** | GAGGCGACCGGTACGATG*CAAACACATGAATGCAGACACCGATTACT*ATA<br>GCAGAGAGATCCTAG | Precursor to no mismatch containing cDNA |
| **CD1.1m** | GAGGCGACCGGTACGATGG*AAACACATGAATGCAGACACCGATTACT*ATA<br>GCAGAGAGATCCTAG | Precursor to 1 nt identical mismatch containing cDNA |
| **CD1.2m** | GAGGCGACCGGTACGATGGT*AACACATGAATGCAGACACCGATTACT*ATA<br>GCAGAGAGATCCTAG | Precursor to 2 nt identical mismatch containing cDNA |
| **CD1.3m** | GAGGCGACCGGTACGATGGTT*ACACATGAATGCAGACACCGATTACT*ATA<br>GCAGAGAGATCCTAG | Precursor to 3 nt identical mismatch containing cDNA |
| **CD1.4m** | GAGGCGACCGGTACGATGGTTT*CACATGAATGCAGACACCGATTACT*ATA<br>GCAGAGAGATCCTAG | Precursor to 4 nt identical mismatch containing cDNA |
| **CD1.5m** | GAGGCGACCGGTACGATGGTTTG*ACATGAATGCAGACACCGATTACT*ATA<br>GCAGAGAGATCCTAG | Precursor to 5 nt identical mismatch containing cDNA |
| **CD1.6m** | GAGGCGACCGGTACGATGGTTTGT*CATGAATGCAGACACCGATTACT*ATA<br>GCAGAGAGATCCTAG | Precursor to 6 nt identical mismatch containing cDNA |
| **CD1.7m** | GAGGCGACCGGTACGATGGTTTGTG*ATGAATGCAGACACCGATTACT*ATA<br>GCAGAGAGATCCTAG | Precursor to 7 nt identical mismatch containing cDNA |
| **CD1.8m** | GAGGCGACCGGTACGATGGTTTGTGT*TGAATGCAGACACCGATTACT*ATA<br>GCAGAGAGATCCTAG | Precursor to 8 nt identical mismatch containing cDNA |
| **CD1.9m** | GAGGCGACCGGTACGATGGTTTGTGTA*GAATGCAGACACCGATTACT*ATA<br>GCAGAGAGATCCTAG | Precursor to 9 nt identical mismatch containing cDNA |
| **CD1.10m** | GAGGCGACCGGTACGATGGTTTGTGTAC*AATGCAGACACCGATTACT*ATA<br>GCAGAGAGATCCTAG | Precursor to 10 nt identical mismatch containing cDNA |
| **CD1.11m** | GAGGCGACCGGTACGATGGTTTGTGTACT*ATGCAGACACCGATTACT*ATA<br>GCAGAGAGATCCTAG | Precursor to 11 nt identical mismatch containing cDNA |
| **CD2.1m** | GAGGCGACCGGTACGATGA*AAACACATGAATGCAGACACCGATTACT*ATA<br>GCAGAGAGATCCTAG | Precursor to 1 nt non-identical mismatch containing cDNA |
| **CD2.2m** | GAGGCGACCGGTACGATGAC*AACACATGAATGCAGACACCGATTACT*ATA<br>GCAGAGAGATCCTAG | Precursor to 2 nt non-identical mismatch containing cDNA |
| **CD2.3m** | GAGGCGACCGGTACGATGACC*ACACATGAATGCAGACACCGATTACT*ATA<br>GCAGAGAGATCCTAG | Precursor to 3 nt non-identical mismatch containing cDNA |
| **CD2.4m** | GAGGCGACCGGTACGATGACCC*CACATGAATGCAGACACCGATTACT*ATA<br>GCAGAGAGATCCTAG | Precursor to 4 nt non-identical mismatch containing cDNA |

| | | |
|---|---|---|
| **CD2.5m** | GAGGCGACCGGTACGATGACCCAACATGAATGCAGACACCGATTACTATAGCAGAGAGATCCTAG | Precursor to 5 nt non-identical mismatch containing cDNA |
| **CD2.6m** | GAGGCGACCGGTACGATGACCCACCATGAATGCAGACACCGATTACTATAGCAGAGAGATCCTAG | Precursor to 6 nt non-identical mismatch containing cDNA |
| **CD2.7m** | GAGGCGACCGGTACGATGACCCTCTATGAATGCAGACACCGATTACTATAGCAGAGAGATCCTAG | Precursor to 7 nt non-identical mismatch containing cDNA |
| **CD2.8m** | GAGGCGACCGGTACGATGACCCTCTCTGAATGCAGACACCGATTACTATAGCAGAGAGATCCTAG | Precursor to 8 nt non-identical mismatch containing cDNA |
| **CD2.9m** | GAGGCGACCGGTACGATGACCCTCTCGGAATGCAGACACCGATTACTATAGCAGAGAGATCCTAG | Precursor to 9 nt non-identical mismatch containing cDNA |
| **CD2.10m** | GAGGCGACCGGTACGATGACCCTCTCGTAATGCAGACACCGATTACTATAGCAGAGAGATCCTAG | Precursor to 10 nt non-identical mismatch containing cDNA |
| **CD2.11m** | GAGGCGACCGGTACGATGACCCTCTCGTCATGCAGACACCGATTACTATAGCAGAGAGATCCTAG | Precursor to 11 nt non-identical mismatch containing cDNA |
| **CD_t-match** | GAGGCGACCGGTACGATGCAAACCTCTTCATGCAGACACCGATTACTATAGCAGAGAGATCCTAG | Precursor to 6 nt mismatch with terminal containing cDNA |
| **CD2.16m** | GAGGCGACCGGTACGATGACCCTCTCGTCCGTACGACACCGATTACTCCATCGCGAGAGATCCTAG | Precursor to 16 nt non-identical mismatch containing cDNA |
| **CD2.20m** | GAGGCGACCGGTACGATGACCCTCTCGTCCGTACTCACCCGATTACTCCATCGCAGAAGGATCCTAG | Precursor to 20 nt non-identical mismatch containing cDNA |
| **Blocker_16** | CAAACACATGAATGCA | 16 nt blocker for CD2.16m |
| **Blocker_20** | CAAACACATGAATGCAGACA | 20 nt blocker for CD2.20m |
| **Primer_PST (phosphorothioated, PST)** | CTT CTG CGA TGG AGT AAT CGG TGT CTG CAT TCA TGT GT*T* T*G | 3′-triple phosphorothioated primer |
| **Primer_nonPST (non-phosphorothioated, non-PST)** | CTT CTG CGA TGG AGT AAT CGG TGT CTG CAT TCA TGT GTT TG | Non- phosphorothioated primer |
| **CD3** | GCT CAA GCA GTT CCT AGC CTC AGA CCT GCG GAT CTG TTA ATT ACG CTA CGG CTG TGT CAC TAA CTA TAC AAC CTA CTA CCT CAT TTC TGC TTG AGC GTC CAG TCC TTT TTG GAC TGG AC | Precursor to alternative cDNA for klenow exo- and phi29 study<br><br>(self-annealing oligonucleotide, does not require a splint) |
| **PST3.0m** | ATT AAC AGA TCC GCA *G *G*T | Phosphorothioated primer for CD3 with 0 nt mismatch |
| **PST3.1m** | ATT AAC AGA TCC GCA *G *G*A | Phosphorothioated primer for CD3 with 1 nt mismatch |
| **PST3.2m** | ATT AAC AGA TCC GCA G *G*A*T | Phosphorothioated primer for CD3 with 2 nt mismatch |

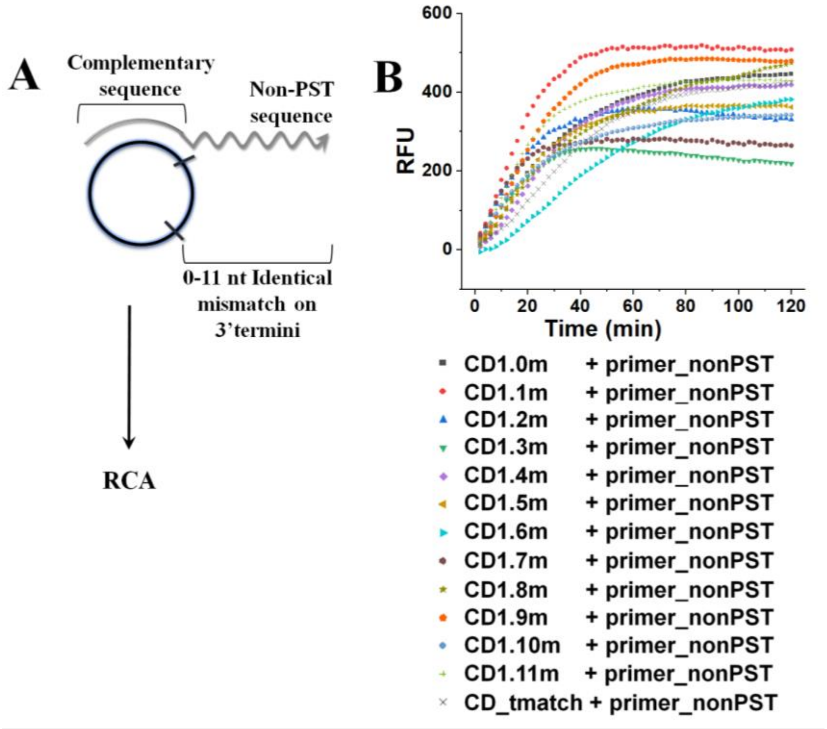


**Figure S1.** Schematics (Panel A) and representative amplification plots (Panel B) involving identical mismatches (0 – 11 nt) in cDNA, primer_nonPST, and phi29 DNA polymerase.

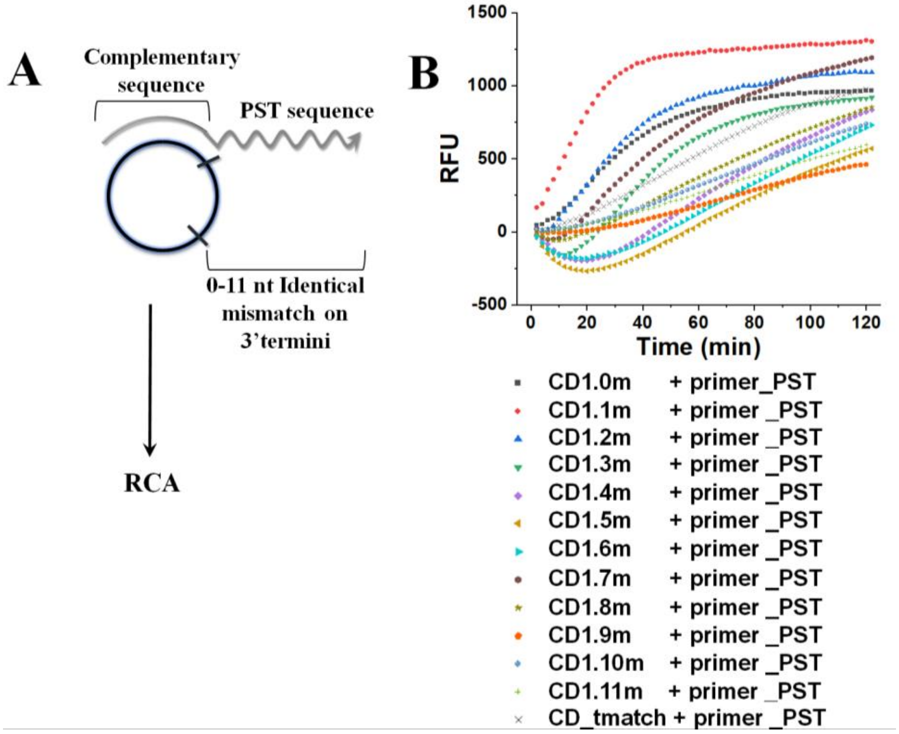


**Figure S2.** Schematics (Panel A) and representative amplification plots (Panel B) involving identical mismatches (0 – 11 nt) in cDNA, primer_PST, and phi29 DNA polymerase.

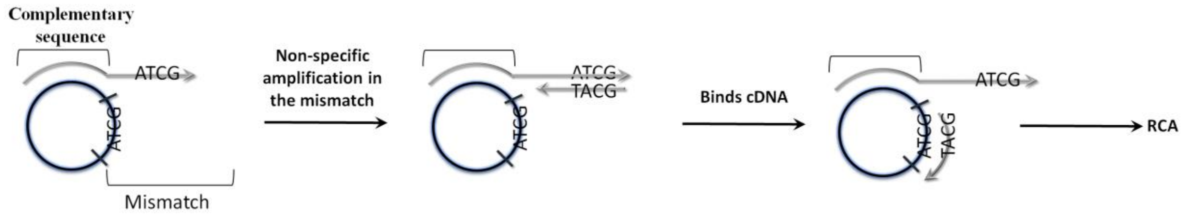


**Scheme S1.** A possible mechanism for terminal mismatch bypass involving non-specific amplification of the identical mismatch region between cDNA and primer_PST.

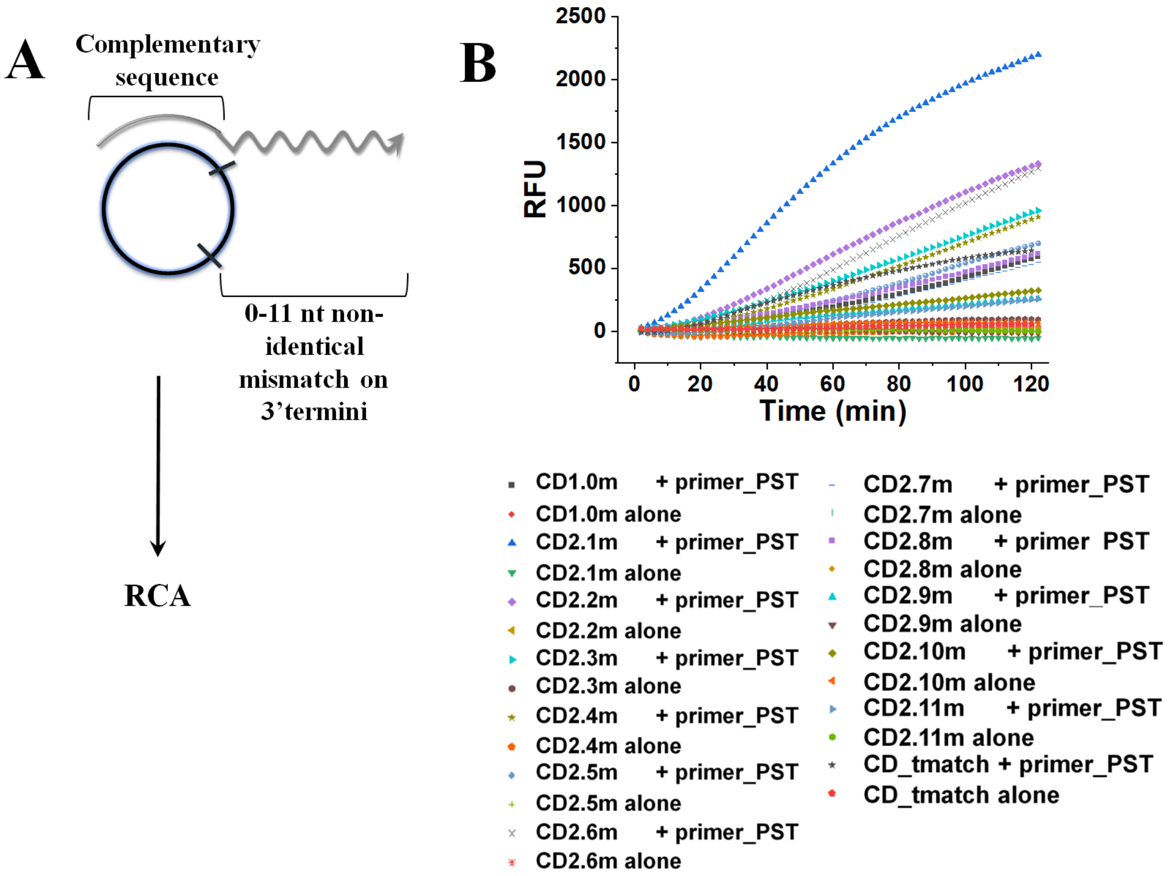


**Figure S3.** Schematics (Panel A) and representative amplification plots (Panel B) involving non-identical mismatches (0 – 11 nt) in cDNA, primer_PST, and phi29 DNA polymerase.

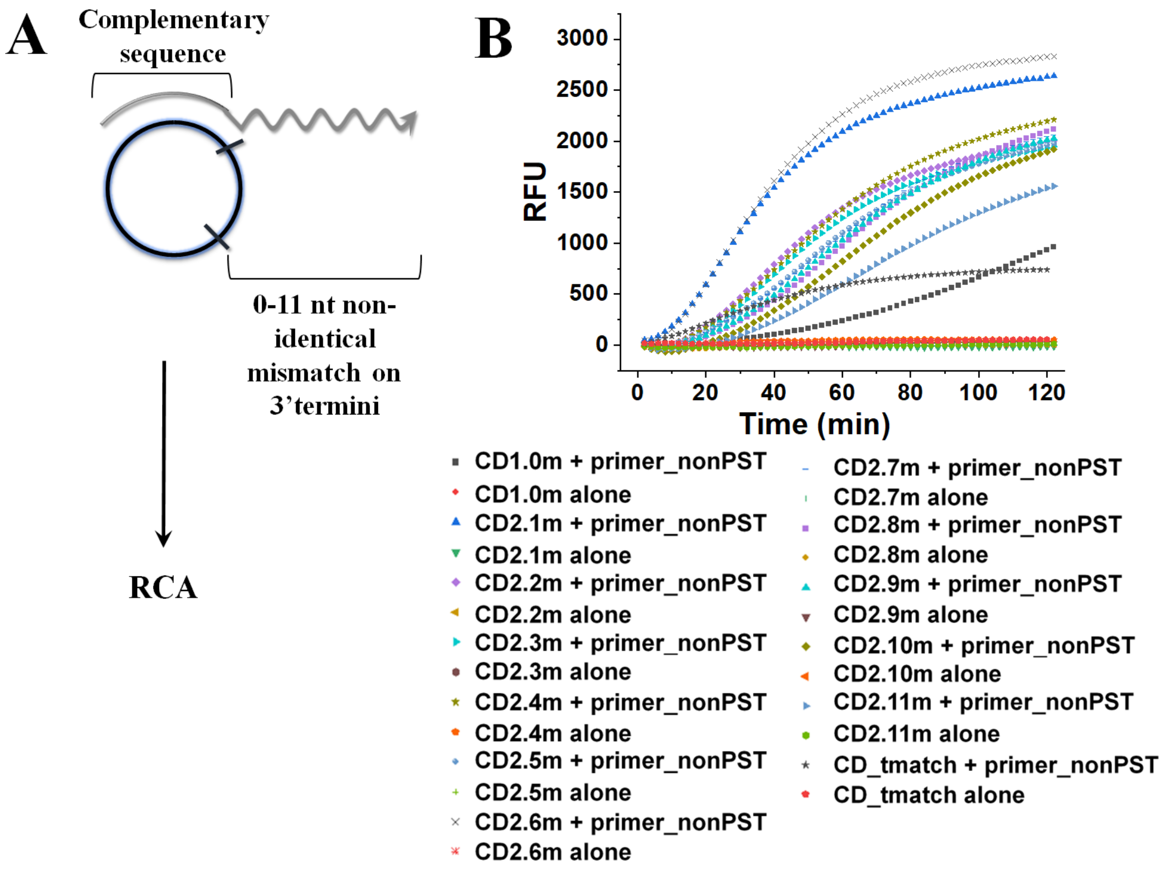


**Figure S4.** Schematics (Panel A) and representative amplification plots (Panel B) involving non-identical mismatches (0 – 11 nt) in cDNA, primer_nonPST, and phi29 DNA polymerase.

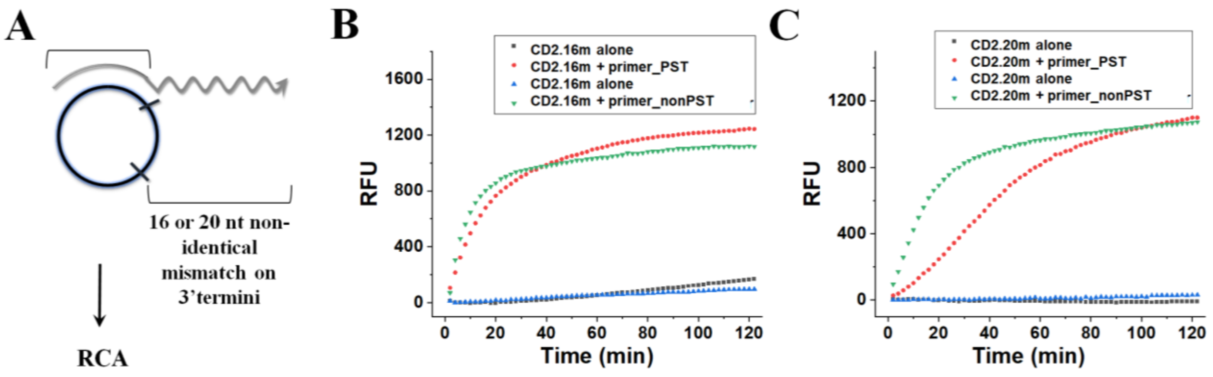


**Figure S5.** Schematics (Panel A) and representative amplification plots (Panel B and C) involving non-identical mismatches 16 and 20 nt, respectively. Phi29 DNA polymerase was used in this study.

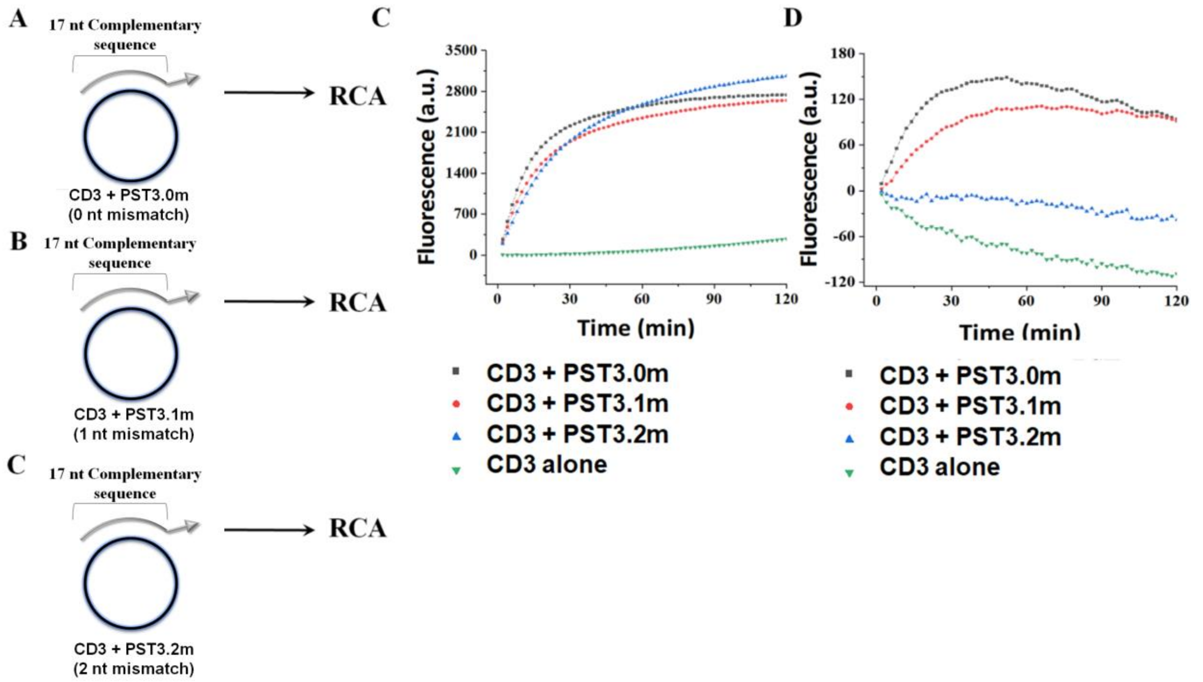


**Figure S6.** Schematics (Panel A – C) of RCA involving self-annealing cDNA CD3 and primers having 0 – 2 nt mismatches. Panel D and E, Amplification involving self-annealing cDNA CD3 and primers having 0 – 2 nt mismatches for phi29 and klenow exo(-) DNA polymerases, respectively.

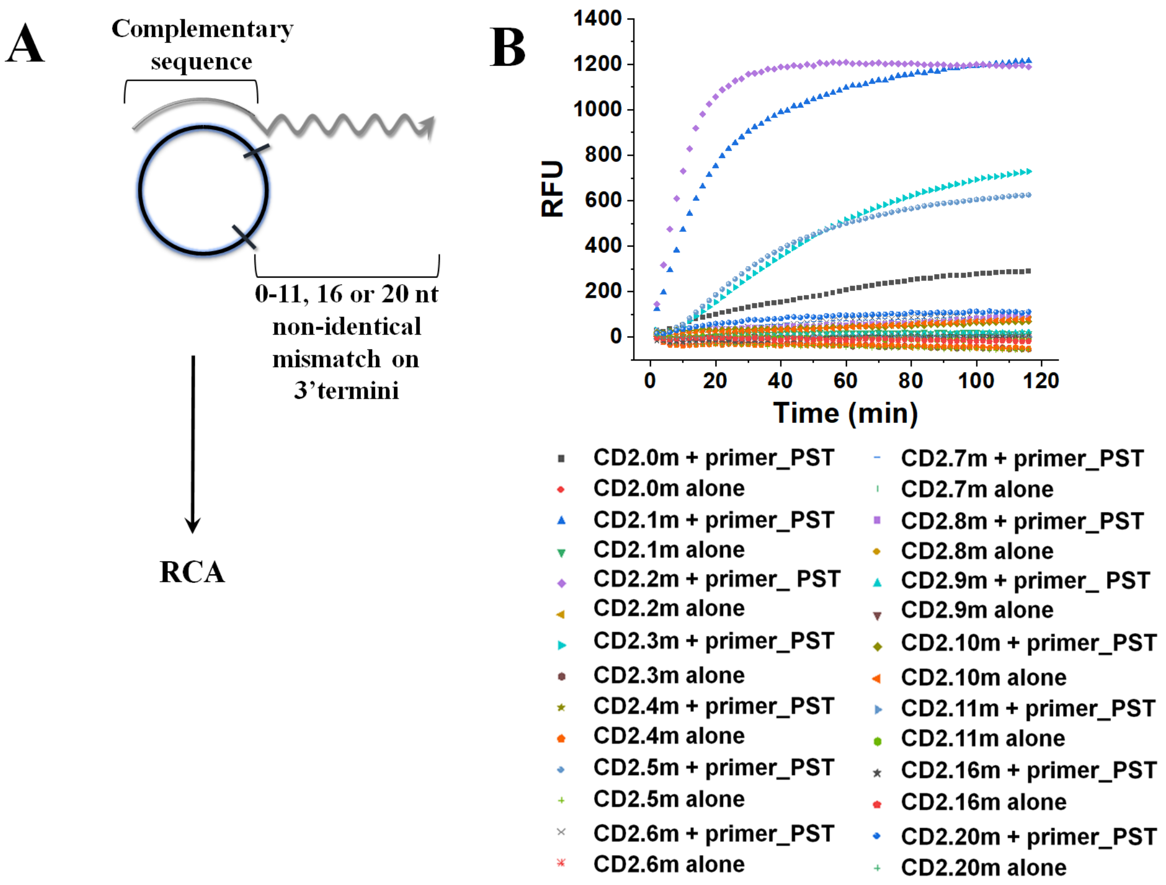


**Figure S7.** Schematics (Panel A) and representative amplification plots (Panel B) involving non-identical mismatches (0 – 11, 16, and 20 nt), primer_PST, and BST large fragment (LF) DNA polymerase.

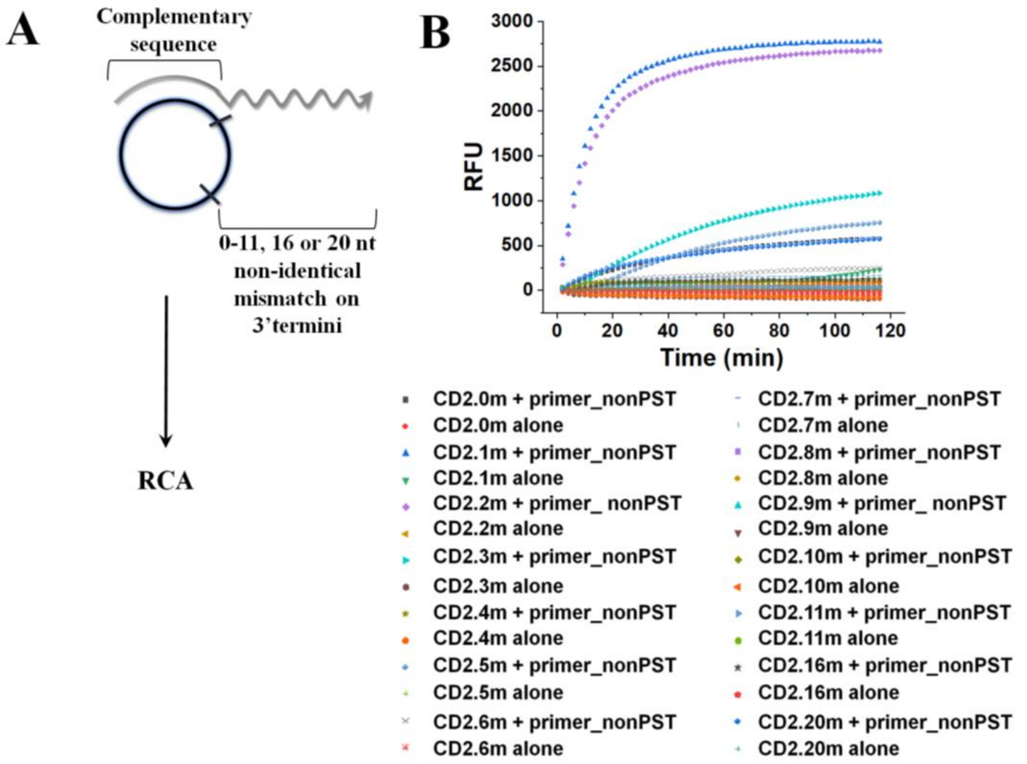


**Figure S8.** Schematics (Panel A) and representative amplification plots (Panel B) involving non-identical mismatches (0 – 11, 16, and 20 nt) in cDNA, primer_nonPST, and BST large fragment (LF) DNA polymerase.

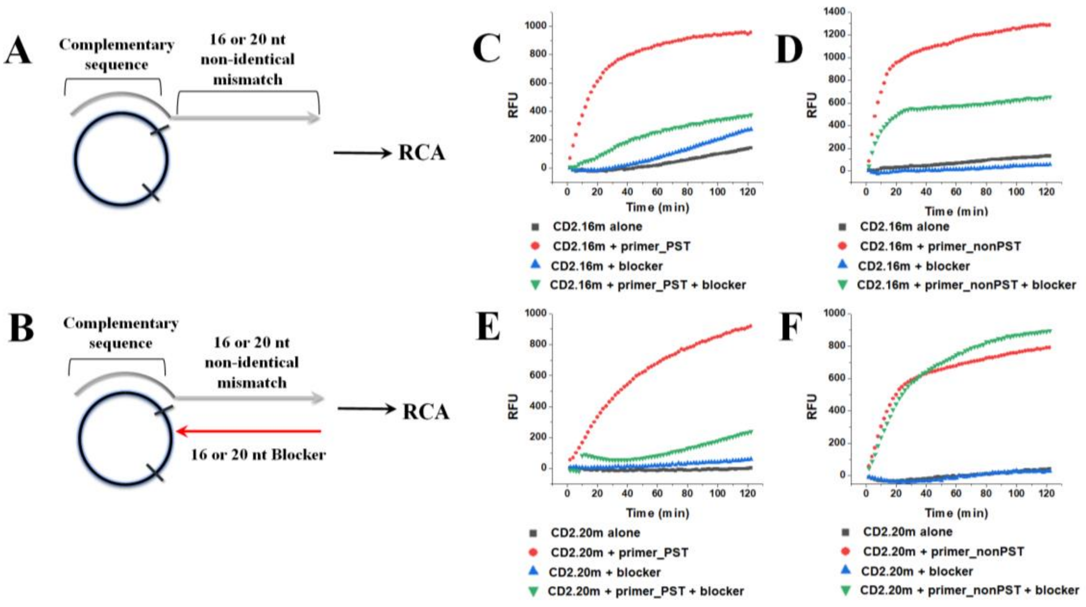


**Figure S9.** Panel A and B, Schematics of "turning on/off" terminal mismatch bypass amplification using a "blocker" sequence. Panel C and D, Representative amplification plots for blocker-mediated regulation of 16 nt terminal mismatch bypass activity for primer_PST and primer_nonPST, respectively. Panel E and F, Representative amplification for blocker-mediated regulation of 20 nt terminal mismatch bypass activity for primer_PST and primer_nonPST, respectively. Phi29 DNA polymerase was used in this experiment.

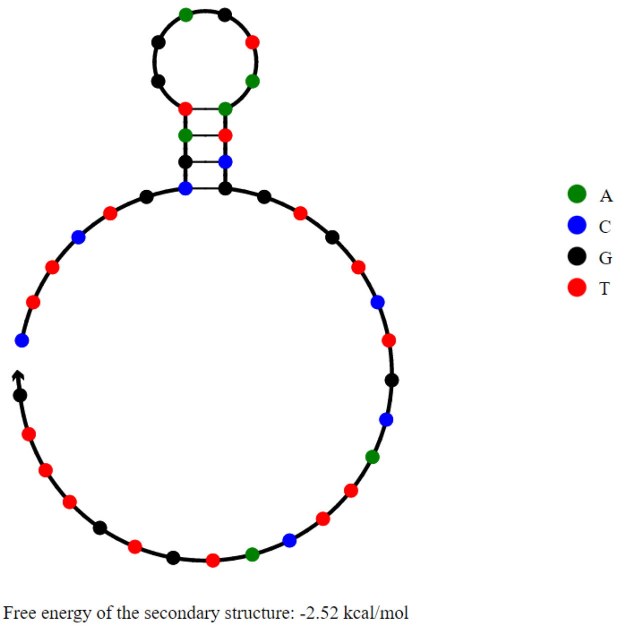


**Figure S10.** NUPACK secondary structures for primer_PST (concentration 1 μM, 1 M $Na^{+}$, 37°C).
